# gcplyr: an R package for microbial growth curve data analysis

**DOI:** 10.1101/2023.04.30.538883

**Authors:** Michael Blazanin

## Abstract

Characterization of microbial growth is of both fundamental and applied interest. Modern platforms can automate collection of high-throughput microbial growth curves, necessitating the development of computational tools to handle and analyze these data to produce insights. To address this need, here I present a newly-developed R package: gcplyr. gcplyr can flexibly import growth curve data in common tabular formats, and reshapes it under a tidy framework that is flexible and extendable, enabling users to design custom analyses or plot data with popular visualization packages. gcplyr can also incorporate metadata and generate or import experimental designs to merge with data. Finally, gcplyr carries out model-free (non-parametric) analyses. These analyses do not require mathematical assumptions about microbial growth dynamics, and gcplyr is able to extract a broad range of important traits, including growth rate, doubling time, lag time, maximum density and carrying capacity, diauxie, area under the curve, extinction time, and more. gcplyr makes scripted analyses of growth curve data in R straightforward, streamlines common data wrangling and analysis steps, and easily integrates with common visualization and statistical analyses.

**Availability:** gcplyr is available from the central CRAN repository (https://CRAN.R-project.org/package=gcplyr), or from GitHub (https://github.com/mikeblazanin/gcplyr).

## Introduction

Characterization of microbial population growth dynamics has been of fundamental and applied interest since nearly the dawn of microbiology [1]. Today, growth curves are a ubiquitous technique to study microbial growth. Indeed, modern automated platforms, including plate readers, can collect high-throughput growth data over time on hundreds of samples simultaneously. Yet, this data-generation capacity has outpaced the development of computational tools to handle and analyze microbial growth data, presenting new challenges.

First and foremost, data are rarely output in the ideal format for analysis, visualization, or publication. Reorganizing data manually can be tedious and fraught with the potential for introduction of errors. Moreover, since output files vary between different plate readers, scripted reorganization may require tailored code for each output format. Despite this, only a few existing software tools provide utilities for streamlined data wrangling and reorganization (Table S1) [2–8].

Once data are reorganized, scientists face the challenge of converting raw data into quantitative microbial traits. Many existing computational tools use parametric analyses of microbial growth curve data (Table S1) [3, 4, 6, 8–19]. Parametric analyses fit a mathematical model of microbial population growth to observed data, then extract the fitted parameter values to quantify traits. This approach can be powerful, but it has drawbacks for many users. First, users must explicitly or implicitly choose a model of growth, making error-prone mathematical assumptions about the form the growth data should take. Moreover, for accurate results users must laboriously verify that their data meet the assumptions of the model and that the fitting process converged on an appropriate solution. Finally, some dynamics do not fit any known models of microbial growth, leaving researchers with few to no options to analyze such growth data.

Given these drawbacks, in recent years some groups have developed software to analyze microbial growth data with non-parametric or model-free approaches (Table S1) [3, 6–8, 14, 15, 17–27]. These analyses make no specific mathematical assumptions about the form of the growth data, instead extracting parameters of interest directly from the data itself or from non-parametrically smoothed transformations of the data. However, many of these tools can only quantify a few traits of interest non-parametrically.

Here, I present my newly-released R package, gcplyr. gcplyr is built as a versatile tool to wrangle plate reader data and carry out model-free analyses. gcplyr is used in R, a popular scripting language for scientific data analysis and visualization. With gcplyr, users can import growth curve data from a wide range of plate reader formats and carry out model-free analyses to extract a number of traits, including lag time, growth rate and doubling time, carrying capacity, area under the curve, and more. Notably, the model-free analyses implemented by gcplyr enable the characterization of dynamics that do not fit traditional mathematical models of microbial growth, and therefore were difficult or impossible to analyze using the parametric methods of many existing tools. Examples of these dynamics include diauxic shifts [28] and growth in the presence of antagonists [e.g. bacteriophages [29]]. Additionally, gcplyr provides a framework for data organization that is flexible and extendable, making it easy for users to run custom analyses or integrate gcplyr workflows with other R packages.

### Implementation

gcplyr is an open-source R package [30] that can be installed with R’s built-in install.packages function from the central CRAN repository (https://CRAN.R-project.org/package=gcplyr), or from GitHub (https://github.com/mikeblazanin/gcplyr). gcplyr has been written to minimize the number of external dependencies needed for installation.

### Usage

gcplyr is usable with a basic working knowledge of the R coding language. A comprehensive manual and tutorials are included with the gcplyr installation and available online (https://mikeblazanin.github.io/gcplyr/). Within R, gcplyr functions make it easy to import data and metadata, merge them with experimental design information, smooth and calculate derivatives as necessary, and analyze curves to produce summary growth curve metrics (Fig 1). gcplyr was designed to easily integrate growth curve analyses with other common R tasks, including plotting with ggplot2 [31], data transformation with tidyverse [32], and statistical analysis.

**Fig 1.**
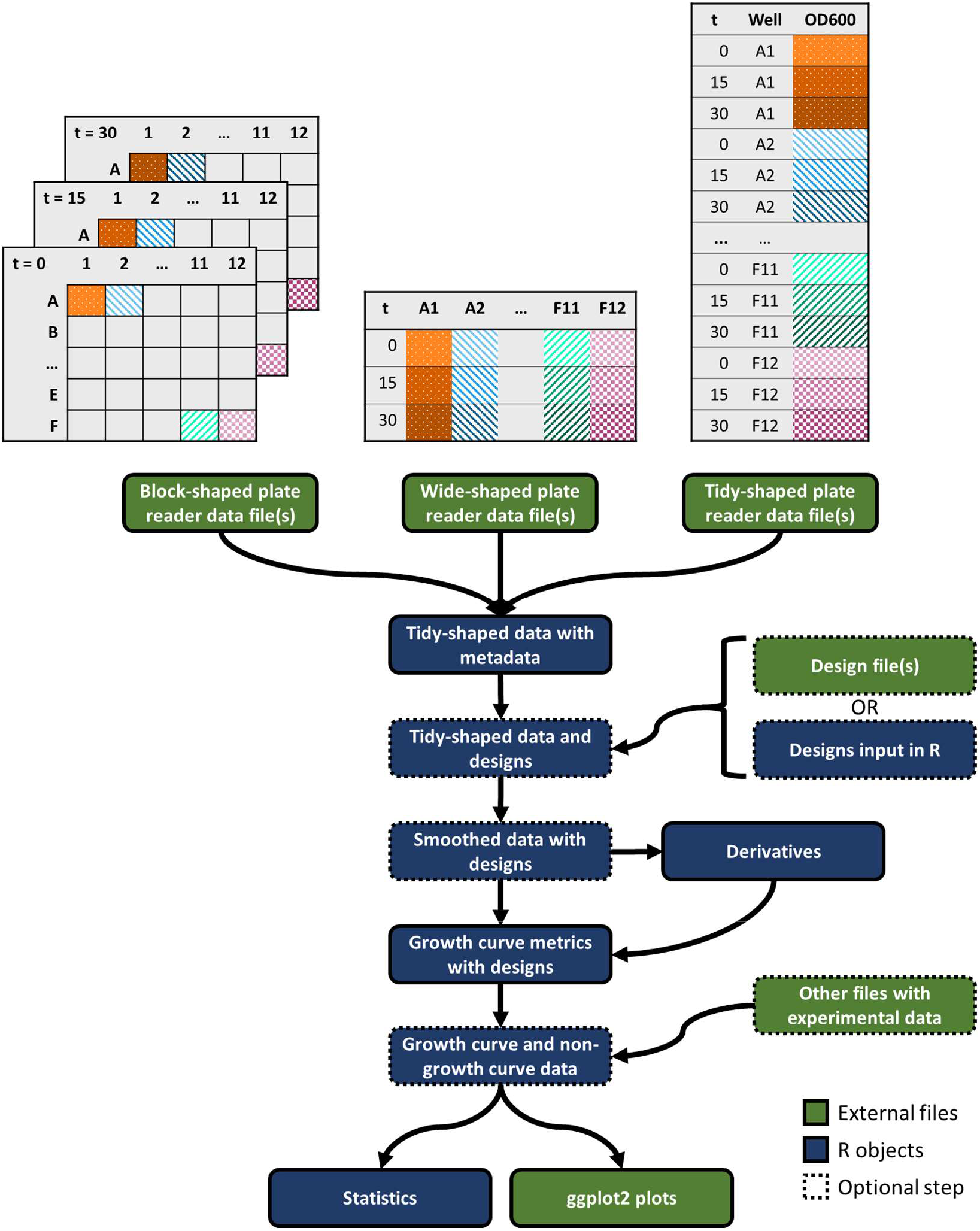
Workflow to use gcplyr to analyze microbial growth curve data. Growth curve data are commonly found in one of three layouts: block-shaped, wide-shaped, or tidy-shaped. Block-shaped data are organized to match the physical layout of the multi-well plate. Wide-shaped data contain one column for each well from a plate. Tidy data contain all the observations in a single column. Gcplyr functions import data files regardless of the data layout, reshape them into tidy-shaped data, then merge them with experimental design information. Data and designs can then be smoothed and have derivatives calculated to extract growth curve metrics. Metrics can be easily merged with other non-growth curve experimental data before statistical analyses and visualization.

#### Inputs

Growth curve data is typically output by plate reader software as block-shaped (also called ‘matrix’) or wide-shaped (also called ‘column-wise’) (Fig 1). gcplyr can import any tabular file containing block-shaped or wide-shaped data, with user-set arguments enabling functionality regardless of where the data is located in the input file. This is an especially useful advance for block-shaped data, where each timepoint is saved separately, and which only one recently-released tool can parse [5]. Additionally, user-set arguments can designate metadata located in input files to be incorporated into the resulting data.

#### Reshaping data

After reading input files into R, gcplyr extends the functionality of tidyr [33] to reshape data into a ‘tidy’ format (also known as ‘long’ format) [34] ideal for subsequent visualization and analysis steps. Tidy data have all observations in a single column, with each unique datapoint with its own row, and additional columns specifying the timepoint, well, and any other information. Tidy data is the best layout for most analyses [34], is consistent with requirements of data repositories like Dryad, and is the expected input for a number of popular R packages [32]. By transforming data into tidy data, gcplyr makes it easy for users to also visualize their data using ggplot2 [31], manipulate their data using dplyr [35], or apply any of the other tidyverse packages to their data [32].

#### Incorporating experimental designs

gcplyr can import experimental design information from user-supplied spreadsheet files, or users can use gcplyr functions to directly generate experimental designs in R. Once imported or generated, these designs can be exported to files or easily merged with imported growth curve data.

#### Smoothing data

The model-free analyses that gcplyr implements can be more sensitive to experimental noise in growth curve data than fitting-based approaches. To address this, gcplyr implements several well-established smoothing functions to smooth noise in raw density data, including moving average, moving median, loess [30, 36–38], GAM [39–42], and smoothing splines. Smoothing functions are tuned by user-set smoothness parameters. Optimal smoothness parameter values can be determined using cross-validation via the caret package [43] to test different parameter values. The package documentation discusses best practices for setting smoothness parameters and smoothing data.

#### Calculating derivatives

Model-free calculation of many growth curve metrics depends on identifying features in the derivatives of density data. gcplyr can calculate both the population rate of growth (derivative) and cellular growth rate (per-capita derivative), using the slope of density over time. However, such derivatives can be sensitive to experimental noise, even if density data has been smoothed. To address this, gcplyr calculates derivatives by fitting a linear regression on a rolling window of multiple points [7], with user-set parameters defining the width of the window. The package documentation discusses best practices for calculating derivatives and setting their smoothing parameters.

#### Characterizing microbial growth

Finally, gcplyr quantifies traits of interest by computing metrics from density and derivatives data. Many of these metrics are quantified by identifying global or local peaks or valleys in the density or derivatives. For local peaks or valleys, gcplyr implements a local extrema-finding algorithm. For instance: maximum growth rate is calculated as the maximum of the cellular growth rate (per-capita derivative); diauxic shifts are identified by calculating the time of a local valley in the population rate of growth (derivative), which identifies a plateau in the original density data; carrying capacity can be calculated as the maximum of density (under the assumption that the population reaches stationary phase during the growth curve). Other metrics are quantified by identifying threshold-crossing events in the density or derivatives. For instance, extinction time can be calculated as the time when density crosses below some threshold value. Finally, some metrics are quantified using purpose-built functions that implement previously-established methods. For instance, area under the curve is calculated by a definite integral of the density data, and lag time is calculated by finding the intersection of the minimum density with a tangent line projected from the point of maximum growth rate [44, 45].

### Brief demonstration of gcplyr analysis of microbial growth curves

Here, as an example, I use gcplyr to quantitatively characterize and compare the growth curves of two bacterial isolates published by [28]: a single ancestral *Pseudomonas fluorescens* clone, and a single experimentally evolved isolate descended from that ancestor. First, I find that the evolved isolate has a lower maximum density (Fig 2A) and area under the curve (Fig 2B) than its ancestor. I then used gcplyr to calculate the cellular growth rate over time from the slope of log-transformed density with a rolling window five data points (75 minutes) wide, finding that the evolved isolate also has a lower maximum growth rate than its ancestor (Fig 2C). All data and code for this analysis are available at https://github.com/mikeblazanin/gcplyr/tree/master/manuscript. Additional examples and in-depth tutorials are included with the gcplyr installation and available online at https://mikeblazanin.github.io/gcplyr/.

**Fig 2.**
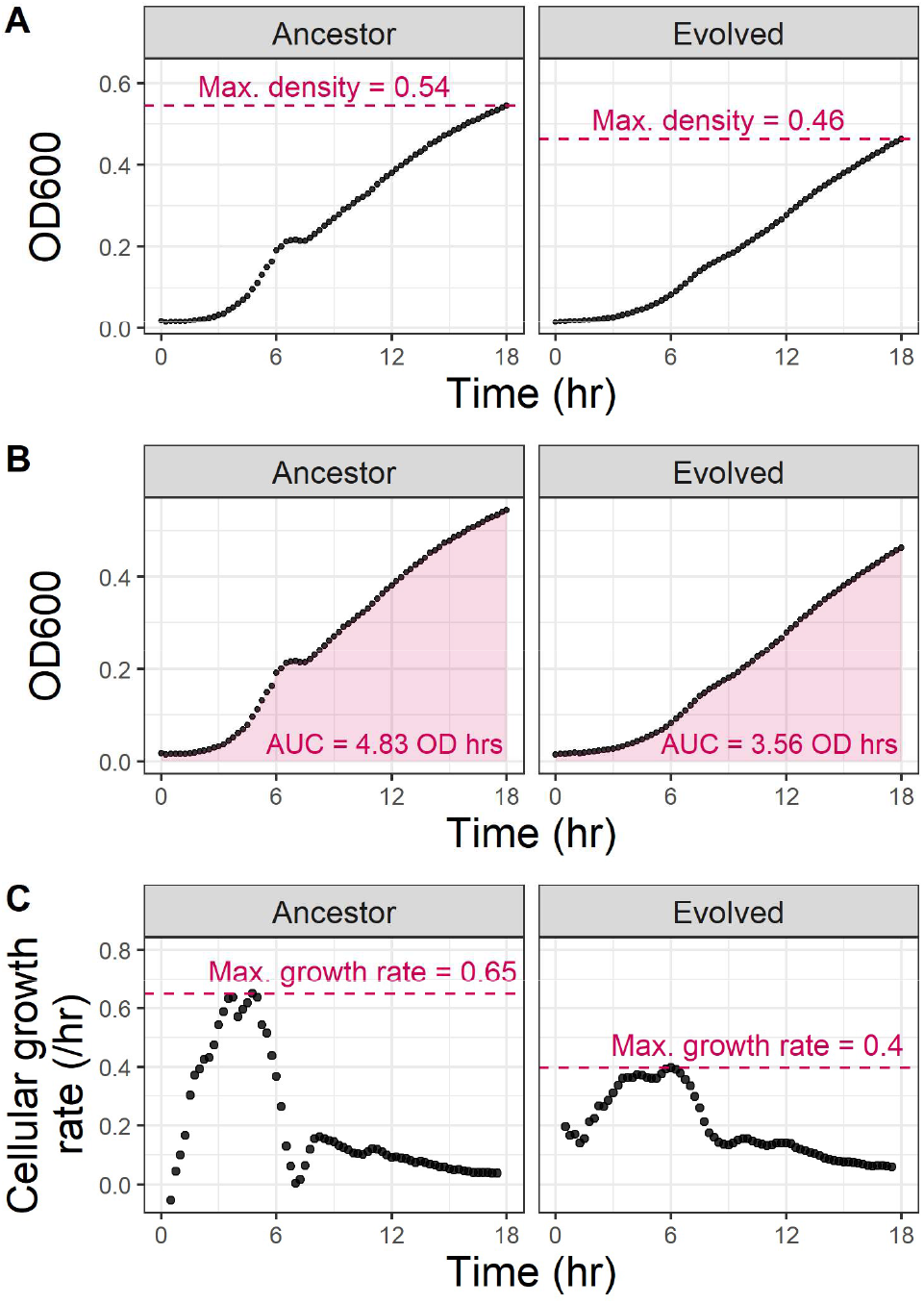
Example analysis of growth curves with gcplyr. I used gcplyr to calculate several common metrics for two previously-published growth curves of *Pseudomonas fluorescens* isolates [28]: one ancestral isolate, and one experimentally evolved descendant isolate. **C**. Here I used gcplyr to calculate cellular growth rate from the slope of log-transformed density with a rolling window five data points (75 minutes) wide.

## Discussion

High-throughput microbial growth data has necessitated computational tools for data wrangling and analysis. Here I introduced gcplyr, a new R package built specifically to address this need.

First, gcplyr wrangles and reshapes growth curve data. gcplyr recognizes that there are common layouts for growth curve data (Fig 1), and it flexibly imports such files. During data reorganization, gcplyr employs a tidy data framework, providing a number of benefits over other data organization approaches:

1. It enables users to integrate as many pieces of experimental design information as desired.
2. It enables users to leverage the general-use functions in gcplyr and other packages to generate custom analyses to identify unique features in their data.
3. It enables users to easily integrate their growth curve analyses with existing visualization (e.g. ggplot2) and statistics packages in R.
4. It enables users to merge growth curve data analyses with other sources of data.

gcplyr is the first package designed for scriptable wrangling of growth curve data in the R programming ecosystem (Table S1). This provides high customizability for users, and enables them to easily incorporate gcplyr with other scripted analyses. In contrast, many existing growth curve analysis tools have little to no data wrangling capability (Table S1), requiring users to reformat files manually. Some packages do provide data reshaping via non-R user interfaces, including bletl via Python scripting [6], and Parsley [5], QurvE [3], and AUDIT [4] via graphical user interfaces.

gcplyr is also the first package in R that is designed for scriptable non-parametric analyses for a wide array of microbial growth traits (Table S1). Many existing tools quantify traits of interest by fitting parametric mathematical models of microbial growth to growth curve data, including biogrowth [16], growthcurver [14] and AUDIT [4]. Some tools provide non-parametric analyses through non-R interfaces, including the Python packages bletl [6], phenom [25], and AMiGA [24], and the graphical user interface-based QurvE [3]. And some scriptable R packages provide non-parametric estimation of specific traits, including area under the curve by growthcurver [14], and maximum growth rate by growthrates [15]. gcplyr builds on these approaches to quantify many different traits non-parametrically, including maximum growth rate, lag time, area under the curve, carrying capacity, diauxie, and more.

gcplyr is a versatile tool designed for users who have a basic level of familiarity with programming in R. Because gcplyr is used via R scripts, it can provide customized data wrangling and growth curve analysis for each user. However, this flexibility comes with a tradeoff: it is less accessible to users who have no experience with R programming, who may be better served by software with graphical user interfaces (GUIs) [3–5, 9–13, 20–23].

All growth curve analyses are limited by their input data. Most frequently, growth curves are collected by automated spectrophotometers, where optical density is used as a proxy for microbial cell density [although see [46] for an alternate proxy for cell density]. However, optical density has limited sensitivity to detect low concentrations of microbial cells, and typically saturates before maximum concentrations of cells are reached [47]. These limitations constrain any analysis of optical density-based growth curves, but may especially affect non-parametric analyses like those of gcplyr.

Additionally, non-parametric analyses have limitations relative to model-fitting approaches. First, input data can contain measurement error, which can more strongly affect non-parametric analyses. To address this, gcplyr includes several smoothing methods, although these are not a panacea for noise. Additionally, non-parametric approaches often require more fine-grained temporal resolution for precise quantification of microbial growth traits. Finally, the traits quantified by non-parametric approaches can sometimes differ from those estimated by model fitting. For instance, non-parametric analyses quantify the maximum realized cellular growth rate, while parametric analysis quantify the intrinsic growth rate [48]. These limitations highlight the relative strengths and weaknesses of the non-parametric approach taken by gcplyr. Parametric methods can be best when density dynamics closely follow known mathematical models of microbial growth and users can validate that model fitting converged on appropriate solutions, while non-parametric analyses enable quantification of traits outside those conditions.

Ultimately, growth curves are a widespread experimental approach in the microbial sciences. However, until now, tools capable of wrangling and doing model-free analysis of growth curve data were limited. By enabling these steps, gcplyr streamlines high-quality model-free analysis for a wide range of applied and fundamental research on microbial growth.

## Acknowledgements

Thanks to Paul Turner for support during this project, to Jyot Antani, Alita Burmeister, Noah Houpt, Albert Vill, and Jordan Lewis for feedback on drafts of this paper, and to all the gcplyr alpha and beta testers.

---

**Table S1.**
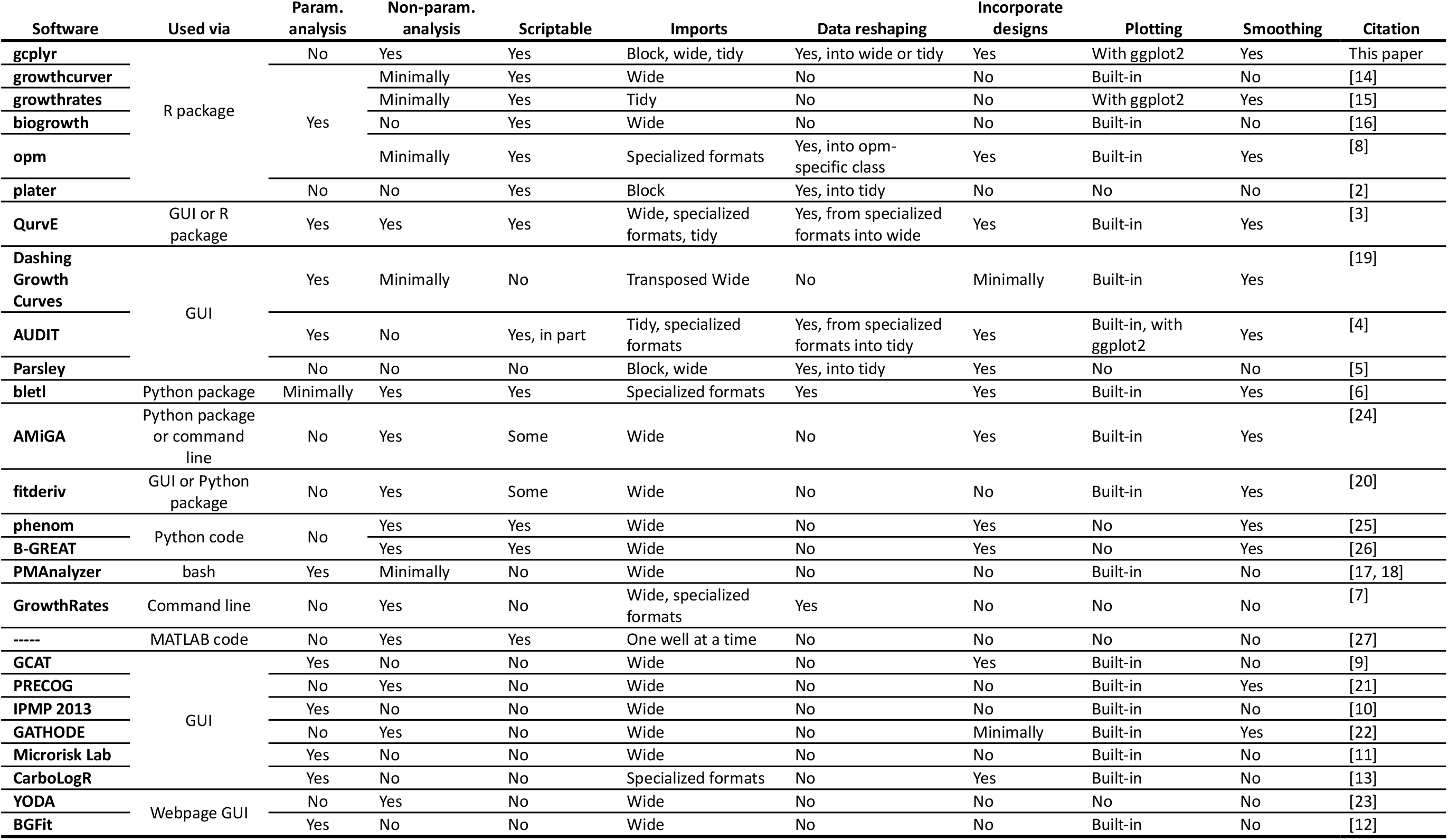
gcplyr and other available microbial growth curve data wrangling and analysis tools.

